# *Escherichia coli* response to subinhibitory concentration of Colistin: Insights from study of membrane dynamics and morphology

**DOI:** 10.1101/2022.01.16.476501

**Authors:** Ilanila Ilangumaran Ponmalar, Jitendriya Swain, Jaydeep K. Basu

## Abstract

Prevalence of wide spread bacterial infections bring forth a critical need in understanding the molecular mechanisms of the antibiotics as well as the bacterial response to those antibiotics. Improper usage of antibiotics, which can be in sub-lethal concentrations is one among the multiple reasons for acquiring antibiotic resistance which makes it vital to understand the bacterial response towards sub-lethal concentrations of antibiotics. In this work, we have used colistin, a well-known membrane active antibiotic used to treat severe bacterial infections and explored the impact of its sub-minimum inhibitory concentration (MIC) on the lipid membrane dynamics and morphological changes of *E. coli*. Upon investigation of live cell membrane properties such as lipid dynamics using fluorescence correlation spectroscopy, we observed that colistin disrupts the lipid membrane at sub-MIC by altering the lipid diffusivity. Interestingly, filamentation-like cell elongation was observed upon colistin treatment which led to further exploration of surface morphology with the help of atomic force spectroscopy. The changes in the surface roughness upon colistin treatment provides additional insight on the colistin-membrane interaction corroborating with the altered lipid diffusion. Although altered lipid dynamics could be attributed to an outcome of lipid rearrangement due to direct disruption by antibiotic molecules on the membrane or an indirect consequence of disruptions in lipid biosynthetic pathways, we were able to ascertain that altered bacterial membrane dynamics is due to direct disruptions. Our results provide a broad overview on the consequence of the cyclic polypeptide, colistin on membrane specific lipid dynamics and morphology of a live Gram-negative bacterial cell.

## 1 Introduction

Traditionally, antibiotics are widely used to treat broad spectrum of prevalent bacterial infections ^1^. Common modes of action of classical antibiotics include targeting the bacterial outer membrane, peptidoglycan as well as the inner membrane and its synthesis machinery ^2^. Additional modes of action include, targeting DNA replication which in turn disrupts bacterial fission and growth, and disrupting specific protein synthesis pathways by suppressing or over-expressing proteins central to the signaling pathways ^3^. Among the above mentioned mechanisms, targeting the bacterial cell envelope is strategically effective and provides specificity against pathogens due to the unique structure and composition of bacterial cell wall ^4^. Therefore understanding the interaction of such membrane targeting antibacterial molecules with the bacterial cell envelope at the molecular level should give us additional insight into antimicrobial action. These studies will enable the development of drug molecules to combat bacterial infections and suppress the emergence of drug-resistant strains ^5^.

Although *Escherichia coli* is a conventional facultative anaerobe that is observed in abundance as a part of gut microbiome, virulent strains are known to cause various infections such as bloody diarrhea, gastroenteritis ^6^. In addition to the diarrheal diseases, the pathogenic *E. coli* also causes meningitis and urinary tract infections ^7^. Enterohemorrhagic *E. coli* are seen in the under-cooked poultry and meat products ^8^. Antibiotics are extensively used in treating multiple bacterial infections in the poultry ^9^ and meat industries ^10^. Uncontrolled usage of antibiotics has led to development of resistant bacterial strains that has necessitated the discovery of a new generation of antibiotics ^11^. Methicillin resistant Staphylococcus aureus (MRSA) and multi-drug-resistant tuberculosis (MDR-TB) are prevalent and challenging to treat with multiple antibiotics ^12,13^. In order to combat these emerging multi-drug resistant strains, colistin, has been re-discovered as a potential drug to be used in combination with other antibiotics ^14^.

Among many antibiotics, colistin (polymyxin E) is a well known membrane active drug which is an amphiphilic positively charged cyclic-polypeptide ^15^. They target the negatively charged lipopolysaccharide (LPS) and replace the Ca^2+^ or Mg^2+^ ions in the core oligosaccharide region. One longstanding hypothesis is that colistin molecules disrupt the physical integrity of the bacterial cell membrane, causing leakage of intracellular contents ^16^. Several recent studies have explored how colistin interacts with the bacterial cell model membranes using transmission electron microscopy (TEM) and other techniques. Membrane permeability due to the action of colistin on small unilamellar vesicles and erythrocytes were shown using calcein leakage and hemolysis assays ^17–19^. Additionally, when colistin attacks the membrane, lipid re-modelling occurs and is reported to form nanoclusters (~2 nm) as well as macroclusters that protrude from the bilayer ^17^. This was confirmed with atomic force microscopy studies. Insights into the binding and penetration of colistin into lipid membrane are investigated using lipid monolayers at air-water interface ^18,20,21^. In contrast to the abundant studies of colistin interacting with LPS in vesicular or monolayer systems, only a few reports of colistin-LPS interactions in a planar bilayer setup have been made. Due to the complex structure of the bacterial cell envelope, constructing realistic bacterial membrane models to carry out in-vitro studies is an active area of research ^22^. Although lipid and protein diffusion on mammalian cell membranes have been discussed in multiple reports, recent experiments have revealed unique protein dynamics on the bacterial cell membrane ^23–25^ which typically occurs on a longer time scale compared to lipid diffusion. However, it is challenging to carry out diffusion measurements using techniques such as fluorescence correlation spectroscopy (FCS) on live bacterial cell membranes due to its small size ^26^. Lipid diffusion is typically measured on model membrane systems ^27,28^ to elucidate the action of membrane active molecules such as anti-microbial peptides, pore-forming toxins ^29–31^, nanoparticles ^32,33^ on the lipid bilayer. The lipid dynamics has revealed significant information about microscopic changes in membrane organization which can correlate with altered cell functionality and signaling and hence is a very crucial probe especially for bacterial cellular response to antibiotics. However, to the best of our knowledge, lipid diffusion on live bacterial cell membranes has been rarely reported till date ^25^.

It is important to note that the colistin concentrations at which bacterial cell lysis are observed are much less in comparison to the reported values used in model lipid membrane ^17,18^. Antibiotics such as colistin are effective only when the concentrations used is higher than the minimum inhibitory concentration (MIC) between consecutive doses prescribed. When the pathogens are treated with sub-MIC antibiotic doses, they tend to evolve resistance against that particular antibiotic through various mechanisms ^34^. Sub-minimum inhibitory concentrations of colistin promotes formation of biofilm in *Acinetobacter baumannii* which makes it more tolerant towards antibiotics ^35^. Laboratory evolution of colistin resistance using sub-MIC of colistin resulted in genetic mutations that helped in retaining the virulent properties of *Acinetobacter baumannii* ^36^. The consequences of sub-MIC colistin exposure on pathogens resulted in providing opportunity for the pathogens to survive in lethal doses. Hence, the main objective of our work is to understand the biophysical as well as morphological changes and response of lipids present in the live cell membrane of the Gram-negative bacterial cell wall towards the accumulation of sub-inhibitory concentration of colistin.

For this study, we have used wild type *E. coli* K12 treated with sub-MIC colistin to understand the biophysical and morphological changes on the cell envelope that occur as a consequence of colistin action. Here, we have successfully applied fluorescence correlation spectroscopy and atomic force microscopy (AFM) techniques to demonstrate that colistin causes significant changes to the morphology and lipid diffusivity on the *E. coli* K12 cell membrane at sub-MIC. Our study has direct implications on identifying how exposure to sub-MIC concentrations of colistin might help in membrane based modifications that could potentially prevent cell lysis. It is imperative to suggest that enhanced lipid dynamics due to membrane reorganisation at sub-MIC is not lethal but makes the bacteria resilient against antibiotics.

## 2 Experimental methods

### 2.1 Materials

*E. coli* NCTC 13846 was procured from Public Health England (PHE) cell cultures in the form of lyophilized pellet which was then reconstituted in the growth media and glycerol stock was prepared for storage purpose. Nutrient media Luria-Bertani (LB), Agar, colistin sulfate, hydrogen peroxide, Nile red was procured from Merck-Sigma Aldrich (India). FM4-64 was procured from Thermofischer Scientific (India).

### 2.2 *E. coli* growth kinetics

Glycerol stock of *E. coli* K 12 was reconstituted in Luria-Bertani (LB) broth and plated in LB agar. The plate was incubated at 37°C overnight and stored at 4°C for further culturing. Primary culture is prepared by incubating a single colony of cells in 5 ml LB and grown overnight at 37°C with continuous shaking. 1% inoculum was used for growing the cells which are subsequently taken for imaging and FCS experiments. For the growth studies, cells from the primary culture were inoculated in 2 ml LB with and without antibiotics (or) H_2_0_2_ in a 24-well plate. The plate was then kept inside a Tecan Plate reader with continuous shaking at 37°C. Turbidity readings were taken at OD_620*nm*_ for every 15 min up to a total duration of 5 hours and the data is fitted to obtain specific growth rates.

### 2.3 *E. coli* membrane labeling and length determination

All the experiments on bacterial membranes were carried out when the cells are within the log phase with the OD of ~0.2 - 0.4. *E. coli* K12 cells that were grown for 2 hours in 5 ml LB at 37°C were labeled with Nile red, rinsed and then fixed to a poly-L-Lysine (PLL) coated glass coverslip. Depending on the experiment, LB media is added with appropriate concentration of antibiotics or H_2_0_2_. PLL helps in fixing the bacteria to the coverslip so that the cells do not move during imaging. 2-5 *μ*l of the re-suspended cells are added to the coverslips and gently spread using a sterile tip. The cells are rinsed with PBS to remove free unbound cells. PBS is then added to the coverslip and during the entire experiment, the cells are immersed in PBS. Images were procured using a SP5-Leica confocal laser scanning microscope (Leica Microsystems, Germany).The length of the bacteria is measured by drawing a line profile along the entirety of the cell and the length measured from multiple bacteria is illustrated in the form of histograms.

### 2.4 Fluorescence Correlation Spectroscopic (FCS) measurements

For all dynamics related experiments, the FCS measurements were carried out on the polar region of the cells and the resulting histograms are the compilation of data from 5-6 different bacteria with approximately 10-20 readings per bacterium. All experiments were performed in triplicate and the mean data are expressed as the mean±standard deviation unless otherwise mentioned. Photon intensities were collected using the Avalanche Photo Diode (APD) detector 581 - 654 nm filter for Nile red and 647 - 703 nm for FM4-64. The intensity was correlated using the PicoQuant Symphotime software. The correlation curves were analysed and fitted using a standard 2D auto-correlation equation. The diffusion coefficient was evaluated using,

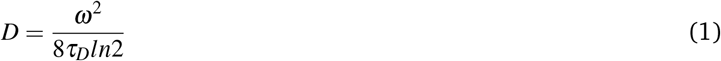

where *ω* is the full width half maximum (FWHM) of the confocal beam ^28^.

### 2.5 Atomic Force Microscopic (AFM) imaging

The bacterial cells are immobilized on the glass coverslips using PLL. The coverslip is placed on the sample holder of AFM stage (Park Systems, South Korea) and the imaging is carried out using tapping non-contact mode in air. Imaging of cells were always performed at room temperature (T=23°C). The images were taken in using PNP-TR2 (NanoWorld, Switzerland) AFM probe with a set point of 0.2 nN force.

## 3 Results and discussion

### 3.1 Role of sub-inhibitory concentration colistin on lipid membrane dynamics of *E. coli*

All the following experiments on bacterial membranes were carried out after 2 hours of incubation when the cells are within the log phase with the OD of ~0.2 (SI Fig.S1). Nile red, lipid-specific hydrophobic dye, was used for the imaging of the live bacterial membrane (Fig. 1 *A,B*). Fig. 1*A,B* images reveal that Nile red is localized along the membrane of bacteria. The control *E. coli* K12 cells are revealed to be rod-shaped with an average length 2.01±0.5 1*μ*m long (Fig. 1*F*), which is typical of a standard *E. coli* ^37^. The mean values are reported with their corresponding standard deviation across three sets of experiments unless otherwise mentioned. Lateral diffusion of Nile red molecules along the lipid bilayer was measured using FCS (see Methods) and is observed as a representative of the lipid lateral diffusion. Molecular dynamics simulation studies on lipid bilayer with Nile red have revealed that the dye prefers to position themselves near the head group of the lipid bilayer ^38^. The histogram for the Nile red diffusion coefficients measured on the wild-type *E. coli* cell membrane is shown in Fig. 1*G*. The average Nile red diffusion is 1.43±0.36 *μ*m^2^/s (Fig. 1*C*). The anomaly parameter, which defines the deviation from free Brownian diffusion, *α* was also fitted and found to be 0.85±0.13 as shown in Fig. 1*H*. The data reveals that the lipid diffusion in bacterial membranes is lower when compared to the model cell membranes where lipid diffusivities on giant unilamellar vesicles ranged from 6 - 12 *μ*m^2^/s^22,39,40^ and supported lipid bilayers ranged from 1 - 6 *μ*m^2^/s^33,41,42^. Such lowered diffusion coefficients are expected in the live bacterial cell membrane and mammalian cell membranes ^43^ which are generally attributed to the complex crowded environment of the cell membrane enriched with various transmembrane proteins. The anomaly parameter, *α*, which is a measure of the deviation from Brownian diffusion also suggests that the presence of membrane proteins in this purportedly crowded environment affects the Nile red diffusion. In order to understand the effects of the membrane disrupting antibiotic, colistin, Nile red diffusion in wild-type *E. coli* is assumed as the control. Thus, any changes in the Nile red diffusion observed in the membrane were solely a consequence of the antibiotic.

**Figure 1.**
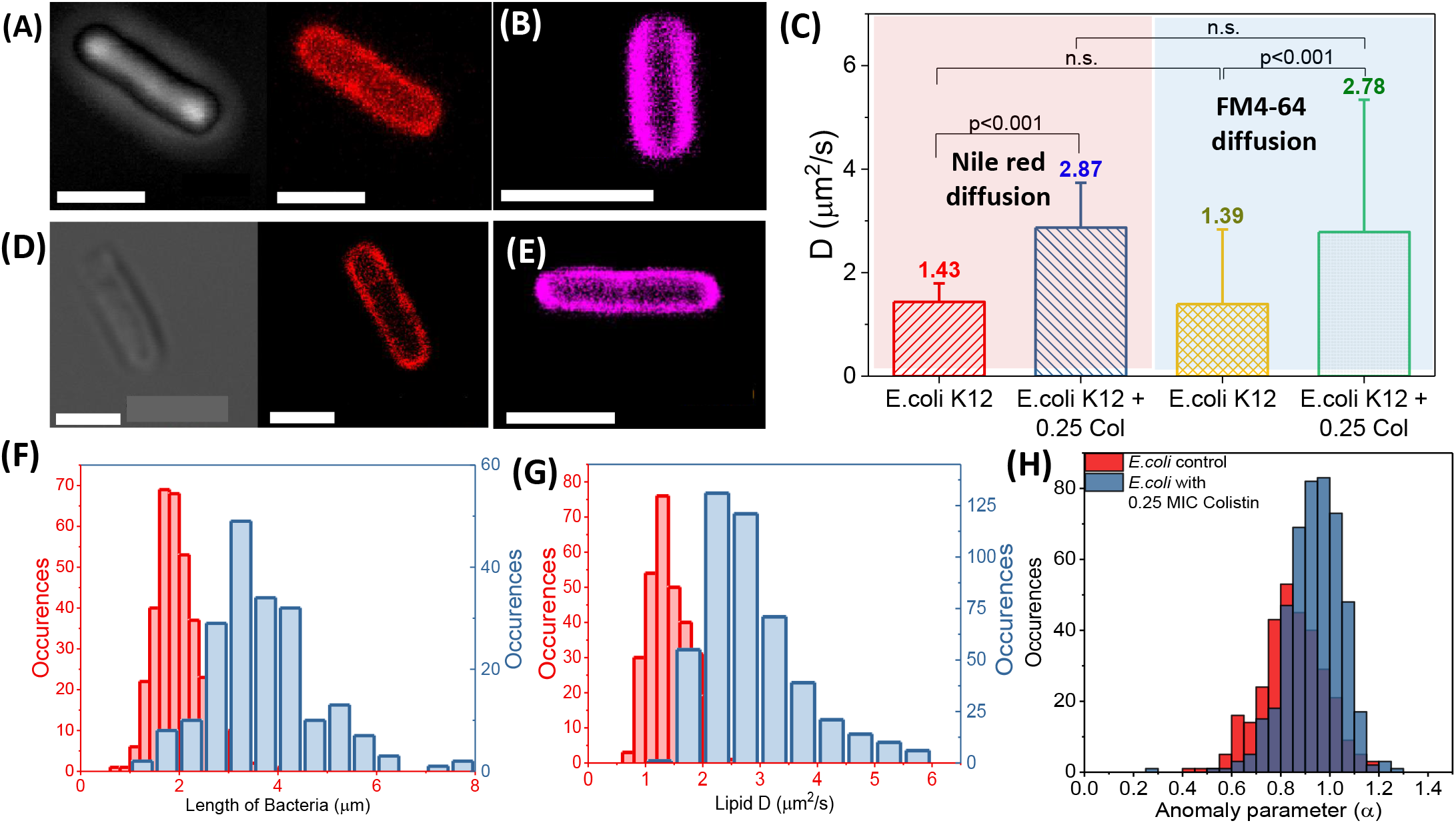
**A.** Phase contrast and confocal images of a typical Nile red labelled *E. coli* wild-type cell. **B** Confocal image of *E. coli* labelled with FM4-64. Corresponding images taken for the cells treated with 0.25 MIC colistin is provided in the panels **D** and **E**. **C.** Average diffusion coefficients measured for Nile red (red) and FM4-64 (blue) labelled cells with and without colistin treatment. **F.** Size distribution of the bacterial cells with and without colistin treatment. **G.**Histogram of Nile red diffusion coefficients in bacterial cells. **H.** Histogram of anomaly parameter (*α*) in bacterial cells. Colour code for panels **F, G, H** - Red bars: Without colistin; Blue bars: With colistin. Scale bar is 2 *μ*m.

Initial experiments are carried out to measure the MIC of colistin on *E. coli* and its corresponding growth kinetics.For all sub-MIC experiments, 0.5 *μ*g/ml concentration of colistin was used which corresponds to 0.25 MIC. We have observed that cells incubated with sub-inhibitory concentration of colistin (0.25 MIC) showed significantly reduced growth rates with a longer lag phase (SI Fig.S1). These results are in good agreement with the earlier reports ^44^. After 2 hours of growth in the presence of 0.25 MIC colistin, the physical attributes such as length of the cells were different from that of the control as shown in Fig. 1*F*. The cells have elongated in size with an average length of 3.68±1.12 *μ*m (Fig. 1*F*). We also observed that a very small population of bacteria had bleb like structures on them (See SI Fig.S4). When the Nile red diffusion was measured, we have also observed that the diffusion coefficients had increased when cells were grown in the presence of 0.25 MIC colistin for two hours. It is to be noted that the Nile red diffusivity is independent of different lengths of bacteria treated with colistin (Fig. 2). The average *D* value was 2.87±0.87 *μ*m^2^/s, which is 2 times greater than that of the control (Fig. 1*C,G*). In contrast to the increase in lipid diffusivities observed on live bacterial cell membranes, recent measurements on model supported lipid bilayers show a decrease in diffusivities when treated with colistin ^45^. The supported lipid bilayers used in these experiments are constituted with phosphatidyl ethanolamine (PE), phosphatidyl glycerol (PG) and cardiolipin (CL) lipids to mimic the inner membranes of Gram-positive and Gram-negative bacterial cell walls. Further, it is interesting to note that their samples were devoid of LPS which are the primary binding sites for colistin ^45^. Interestingly, another report on molecular dynamics simulation have shown that the antibiotic, polymyxin B, which is in the same class of molecules as colistin, is reported to loosen the outer membrane and stiffen the inner membrane ^46^.

**Figure 2.**
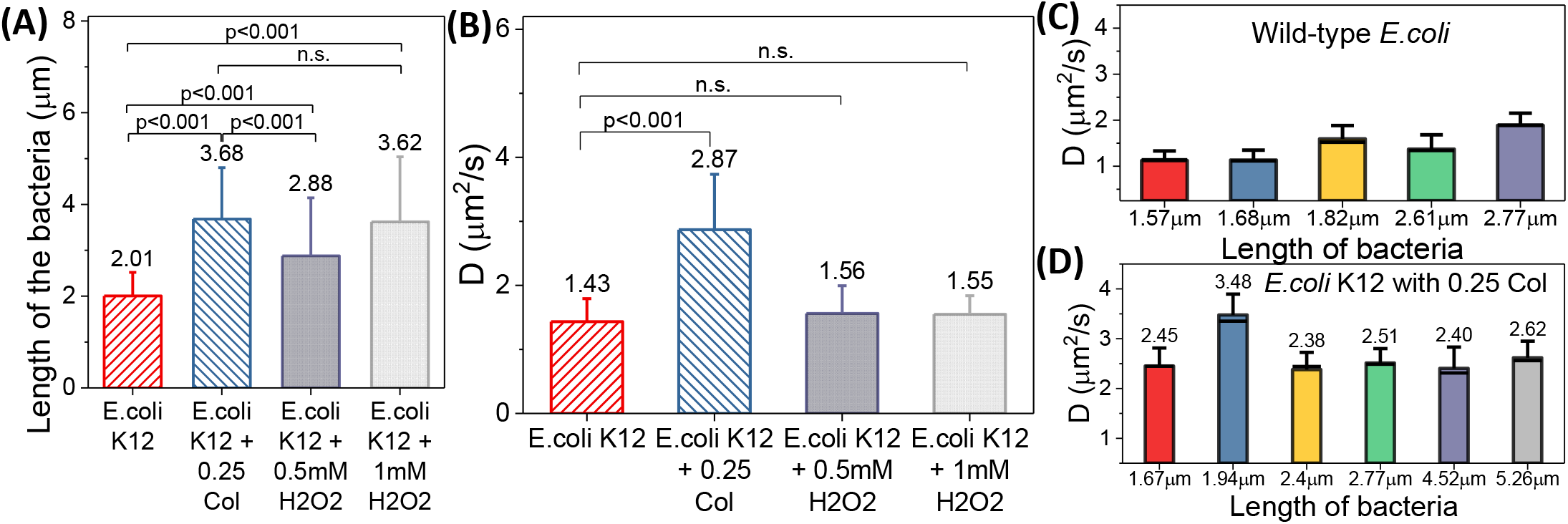
Comparison of average bacterial length (**A**) and Nile red diffusion coefficients (**B**) for wild-type *E. coli* with colistin and hydrogen peroxide treated cells. Colour code: Red - Wild-type *E. coli*; Blue - Colistin treated cells; Purple - 0.5mM H_2_O_2_ treated cells; Grey - 0.5mM H_2_O_2_ treated cells. Average Nile red diffusion coefficients of single wild-type *E. coli* (**C**) and colistin treated *E. coli* (**D**) across different cell lengths.

**Figure 3.**
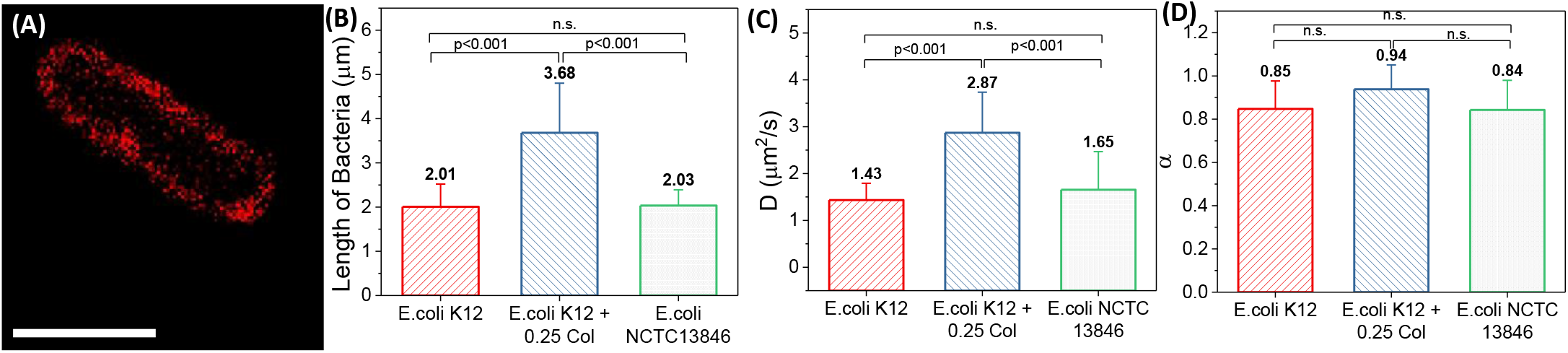
**A.** Confocal fluorescent image of a sample *E. coli* NCTC 13846 cell labelled with Nile red. Comparison of wild-type *E. coli* average cell length (**B**), Nile red diffusion coefficients (**C**), and anomaly parameter *α* (**D**) with 0.25 MIC colistin treated cells and *E. coli* NCTC 13846 grown in 10 *μ*g/ml colistin. Scale bar is 2 *μ*m.

Comparing the diffusion coefficients measured on the wild-type and antibiotic treated *E. coli* cells as given in Fig. 1*C*, we can see that there is significant increase observed in the colistin treated cells. Focusing on mean values of the anomaly parameters *α* extracted from these membrane dynamics for the antibiotic treated cells (Fig. 1*H*), the values did not significantly differ from the wild-type cells demonstrating the anomalous behaviour of the lipid dynamics observed even during the colistin interaction. Although it is well known from the plasma membrane literature that an increase in lipid diffusivity can be attributed to either disordering of the membrane? ^47^ or an increase in lipid free area? ^48^ measurements of lipid dynamics in bacterial membranes exposed to antibiotics are lacking in the literature. Hence, some of the potential possibilities that can cause such enhanced diffusion coefficients include changes in the Nile red itself due to the action of colistin, the physical elongation of the cell, or changes in the lipid membrane due to direct interaction of the colistin molecule on the membrane. Additionally, in the case of live bacterial cells, one of the important factors that can potentially contribute to changes in Nile red diffusion are variations in membrane composition that are observed when *E. coli* adapts to acquire antibiotic resistance ^49^. In the following sections, we have systematically verified the above mentioned possibilities using different methodologies to identify the exact reason behind enhanced lipid fluidity due to colistin interaction at sub-MIC levels.

Further to prove that our above mentioned observation is independent of the lipid dye used for the measurements, we have used another well known lipophilic styryl dye, FM4-64 (N-(3-Triethylammoniumpropyl)-4-(6-(4-(Diethylamino) Phenyl) Hexatrienyl) Pyridinium Dibromide) that has been used widely for imaging cell membranes ^50^. From Fig. 1**B, E**, we have observed that the FM4-64 dye was spread homogeneously on the *E. coli* cell envelope although its specificity in binding to the particular component of the cell envelope is not yet reported ^50^. When FCS was carried out on the FM4-64 labelled wild-type cells that are treated and untreated with colistin, the diffusion values measured were similar to the Nile red diffusion as shown in Fig. 1*C*. The FM4-64 diffusion coefficient of the wild-type cells are observed to be 1.39±1.04 and for colistin treated cells, it is reported to be 2.78±1.55. This gives us additional evidence that the Nile red diffusion coefficients are indeed representative values of lipid diffusion on the cell membrane.

### 3.2 Lipid dynamics do not depend on cell elongation and lipid composition

In order to correlate the elongation caused due to antibiotic stress on bacteria with lipid diffusion, we carried out experiments by incubating the cells with hydrogen peroxide to induce oxidative stress. It has been reported earlier that the oxidative stress induced by H_2_O_2_ affects the cell morphology and protein synthesis ^51^. Hence, we incubated the cells with 0.5 and 1 mM H_2_O_2_ for a period of two hours and repeated the imaging, FCS and growth experiments. Size analysis on the H_2_O_2_ treated cells showed that it does affect the cell morphology and the average length of the bacteria is found to be 3.62±1.42 *μ*m for (1 mM H_2_O_2_) which is significantly larger compared to the control cells and is similar to the colistin treated cells. Concentration dependent size distribution is provided in Fig. 2*A*. These sizes are comparable to the colistin treated cells. When Nile red labeled cells were observed under confocal beam for FCS, the average lipid diffusivity was 1.55±0.29 *μ*m^2^/s with an average *α* values of 0.90±0.14. The Nile red diffusion values shown in Fig. 2*B* shows that the diffusivity of lipids were not enhanced as observed in the colistin treated cells. This particular experiment was carried out for 0.5 mM H_2_O_2_ and the lipid diffusivities were not significantly different when compared with Nile red diffusion measured on the control cells.

Additionally to validate our earlier claim that the cell growth rates do not affect lipid diffusivities, the growth curves were evaluated for the cells incubated with 0.5 and 1 mM H_2_O_2_ based on the time-dependent turbidity values measured at 620 nm. As provided in SI Fig.S1, growth rates decreased for cells under oxidative stress and the lag time increased with increasing concentration of H_2_O_2_. This retarded growth was similar to trends observed with colistin treated cells (SI Fig.S1) where we found that the lipid diffusivities were significantly enhanced from the control experiments (Table. 1).

**Table 1.** Physical properties of the wild-type and treated *E. coli* cells

| 2*Sample | Length of the cell ($\mu\text{m}$) | | Lipid D ($\mu\text{m}^2/\text{s}$) | | Anomaly parameter ($\alpha$) | |
| --- | --- | --- | --- | --- | --- | --- |
|  | Mean | SD | Mean | SD | Mean | SD |
| Control | 2.01 | 0.51 | 1.43 | 0.36 | 0.85 | 0.13 |
| 0.25 MIC colistin | 3.68 | 1.12 | 2.87 | 0.86 | 0.94 | 0.11 |
| 0.5 mM $\text{H}_2\text{O}_2$ | 2.88 | 1.27 | 1.56 | 0.43 | 0.87 | 0.13 |
| 1 mM $\text{H}_2\text{O}_2$ | 3.62 | 1.42 | 1.55 | 0.29 | 0.90 | 0.14 |
| *E. coli* NCTC 13846 | 2.03 | 0.36 | 1.63 | 0.81 | 0.84 | 0.14 |

Similarly, when compared with the control cells where the cells were grown in the absence of stress, the growth rates for cells treated with H_2_O_2_ were distinctly slower. However despite these slower growth rates, the lipid diffusivities were similar to the control cells (Table. 1) and independent of the concentration of H_2_O_2_. The extent of elongation with the H_2_O_2_ treated cells were also similar to the elongation observed with the colistin treatments (Table. 1), providing additional confirmation to the lack of correlation between increased lipid diffusion and cell elongation. Taken together, the results observed on cells treated with H_2_O_2_ and cells treated with colistin provide strong evidence for the following. The increase in lipid diffusion depends on the nature of the stress (oxidative or antibiotic) and does not depend on the cell growth kinetics or the extent of elongation. These results suggest that the observed changes in lipid diffusion arise from a modulation in the membrane lipid composition at sub-inhibitory levels of antibiotics.

We have further extended our lipid dynamics experiments in another strain of *E. coli*, which has a gene that introduces modifications on the outer leaflet of the cell envelope by introducing the phosphatidylethanolamine groups. This strain NCTC 13846 (National Collection of Type Cultures) has a plasmid that encodes mcr-1 (mobilized colistin resistance-1) gene that has been reported to modify the charge on the membrane by introducing phosphatidylethanolamine residue (pEtN) on the head group of lipid A thereby neutralising the negative charge of LPS ^49^. We wanted to explore the relationship between the enhanced lipid diffusion observed at sub-MIC of colistin to a different strain of *E. coli* to check if there is any change in lipid composition of the cell envelope that is causing the lipid disorderness and surface roughness. The *E. coli* NCTC 13846 strain was grown in the presence of 10 *μ*g/ml of colistin. Although the lipid compositions of the mutant strain is modified in comparison to the wild-type *E. coli*, interestingly, we could observe that these strains are similar to wild-type *E. coli* cells in the absence of colistin and are significantly different from the colistin treated wild-type cells. The *E. coli* NCTC 13846 strain grown in the presence of colistin was 2.03±0.36 *μ*m in length with the mean Nile red diffusion coefficient of 1.63±0.81 *μ*m^2^/s which is significantly different from the colistin treated wild-type *E. coli* cells indicating that lipid modifications are not related to the changes in the membrane fluidity and incubation of cells with sub-MIC colistin does not cause lipid alterations similar to mcr-1 gene based lipid modifications.

### 3.3 Role of sub-inhibitory concentration of colistin on surface morphology of the *E. coli* cells

The surface morphology of the cells are studied using the standard atomic force microscopic imaging in air. The control cells as well as colistin treated cells are imaged in air using non-contact mode which is typically carried out for imaging bacterial cells ^52^. The cells were imaged after ~ 5-10 min exposure to air for all the samples to avoid imaging artifacts due to drying. While imaging in air, we could observe that upon treatment with colistin, the physical attributes of the *E. coli* such as the cell width and the cell height were ~ 4 *μ*m and ~ 500 nm (Fig. 4*D-F*). In comparison to the wild-type cells that are ~ 2 *μ*m wide and ~ 300 nm high (Fig. 4*A-E*), the colistin treated cells are larger in physical morphology. Although, we did not observe such variation in width while imaging in confocal fluorescence microscope due to resolution limitation, one of the reasons that can attribute to such changes is that the cells are imaged in air during AFM. Such morphological differences between air-imaging and liquid-imaging was reported in earlier studies ^52^.

**Figure 4.**
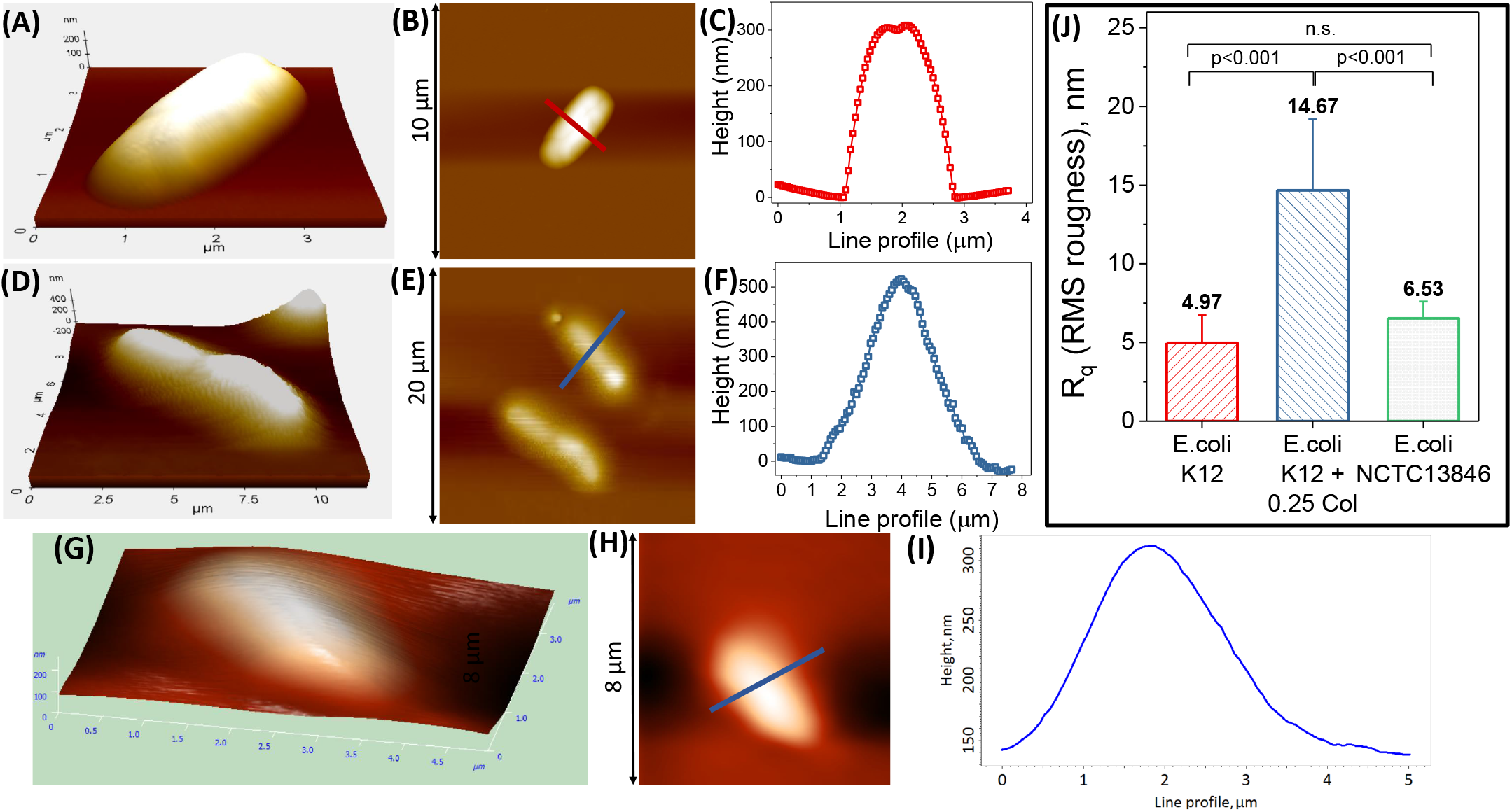
Atomic force microscopic image of wild-type *E. coli* in 3-dimension (**A**) and 2-dimension (**B**) taken in air and corresponding line profile across the cell is plotted in panel C. Similar images and line profiles are plotted for wild-type *E. coli* treated with 0.25 MIC colistin (**D, E** and **F**) and *E. coli* NCTC 13846 strain (**G, H** and **I**). **J.** Mean surface roughness measured across different cells.

While colistin treated wild-type cells were morphologically different, the mutated strain *E. coli* NCTC 13846 was on average ~ 2.5 *μ*m wide and ~ 150 nm high (Fig. 4*G-I*), structurally similar to the wild-type *E. coli*. In order to observe colistin interaction with the surface morphology, additional analysis was carried out to measure the root-mean squared roughness denoted by R_*q*_. The R_*q*_ was 4.97±1.75 nm, 14.67±4.5 nm and 6.53±1.37 nm for the wild-type *E. coli*, colistin treated *E. coli* as well as *E. coli* NCTC 13846 respectively (Fig. 4*J*). The roughness was significantly larger for the colistin treated *E. coli* in comparison to the wild-type cells and *E. coli* NCTC 13846. Our results on live *E. coli* cells are similar to the reported roughness values observed on *Acinetobacter baumannii* ^53^, where the authors reported an increase in the surface roughness as a function of colistin concentration. Increased roughness is a consequence observed due to the direct interaction of sub-MIC colistin on the cell membrane causing membrane disorder and rearrangement which is the major cause for enhanced lipid diffusion coefficients. Correlating the enhanced dynamics measured using FCS and the modified membrane roughness observed from AFM images, we can establish the effect of sub-MIC concentration of colistin on live bacterial membranes.

## 4 Conclusions

In this work, we have systematically investigated the changes in lipid diffusion in live bacterial cell membranes exposed to sub inhibitory concentration of colistin ^14^. Upon treating the *E. coli* K12 bacterial cells with 0.25 MIC colistin for two hours, we observed elongated cells with a factor of two increase in the lipid diffusion coefficients. It is imperative to note that lipid dynamics observed from inner membrane preferring FM4-64 dye ^54^ indicated the altered lipid properties in both inner as well as outer membrane of the *E. coli*. Studying the systematic partitioning of the diffusion coefficients across the bacterial lengths in each system and across the different systems, suggests that the extent of elongation are uncorrelated with the changes observed in the lipid dynamics.

We were able to eliminate the possibility of correlation between growth kinetics, cell elongation and lipid dynamics, which left us with two other possibilities to explain the observed changes in lipid diffusion. The change in diffusivities could be attributed to an outcome of direct interaction between antibiotic molecules on the membrane or an indirect consequence of disruptions in lipid biosynthetic pathways. During the elongation process, it might be possible that the lipid composition is also modulated resulting in the observed increase in diffusivities. Our results from colistin treated *E. coli* NCTC 13846 helps reveal that the altered lipid dynamics observed is due to direct interaction of the colistin molecules on the live inner and outer cell membranes. Finally, we conclude that the dynamics of both inner and outer membrane lipids plays a significant role in the action of a membrane targeting antibiotic at sub-MIC concentration which might have implications in the resistance evolution.

## Author Contributions

IIP, JS performed the experiments and analysed the data. IIP, JS, JKB wrote the paper.

## Acknowledgements

The authors thank Prof. K. Ganapathy Ayappa for his constructive guidance during experiments and Prof. Sandhya Visweswaraiah for her feedback that helped improve the manuscript. The authors thank financial support from Institute of Science,(IISc), Bengaluru, through the ministry of human resources development (MHRD) funded institute of eminence (IOE) project.

## Supplemental Information

**Figure S1:**
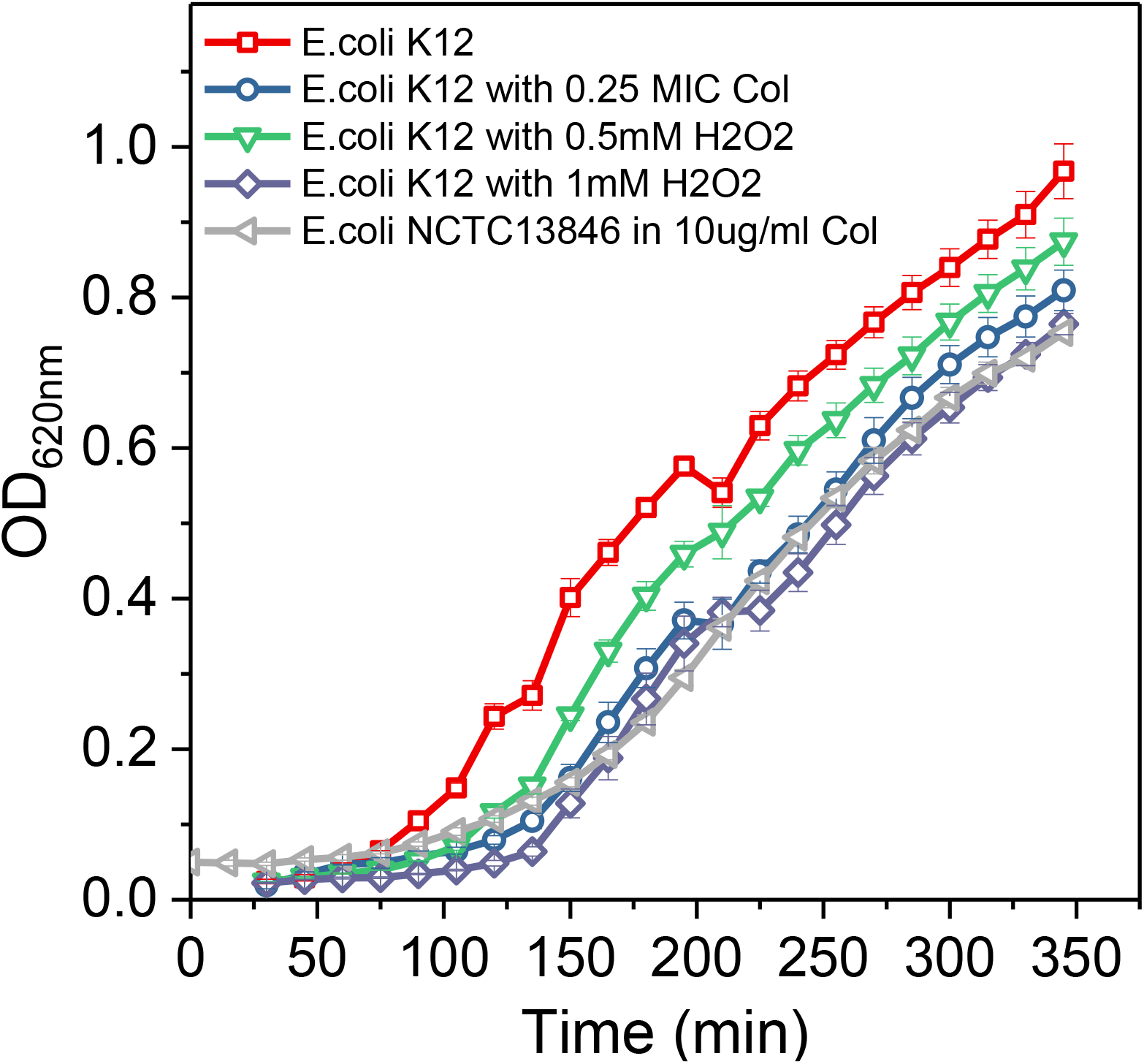
Growth curve of *E. coli*: Absorbance measurements at OD_620*nm*_ was monitored as a measure of the bacterial cell growth. 1% inoculum of *E. coli* K12 as well as *E. coli* NCTC13846 was added to the growth medium and incubated at 37 °C with continuous shaking. The data was collected at a time interval of 15 min. Growth curves of *E. coli* K12 subjected to colistin treatment (blue circles) as well as hydrogen peroxide treatment (green triangles and purple diamonds) indicate slower growth rate in comparison to the wildtype *E.coli* K12 (red squares). Interestingly, the mutant strain, *E. coli* NCTC13846 (grey triangles) grown in the presence of 10*μ*g/ml colistin also indicated a slower growth rate

**Figure S2:**
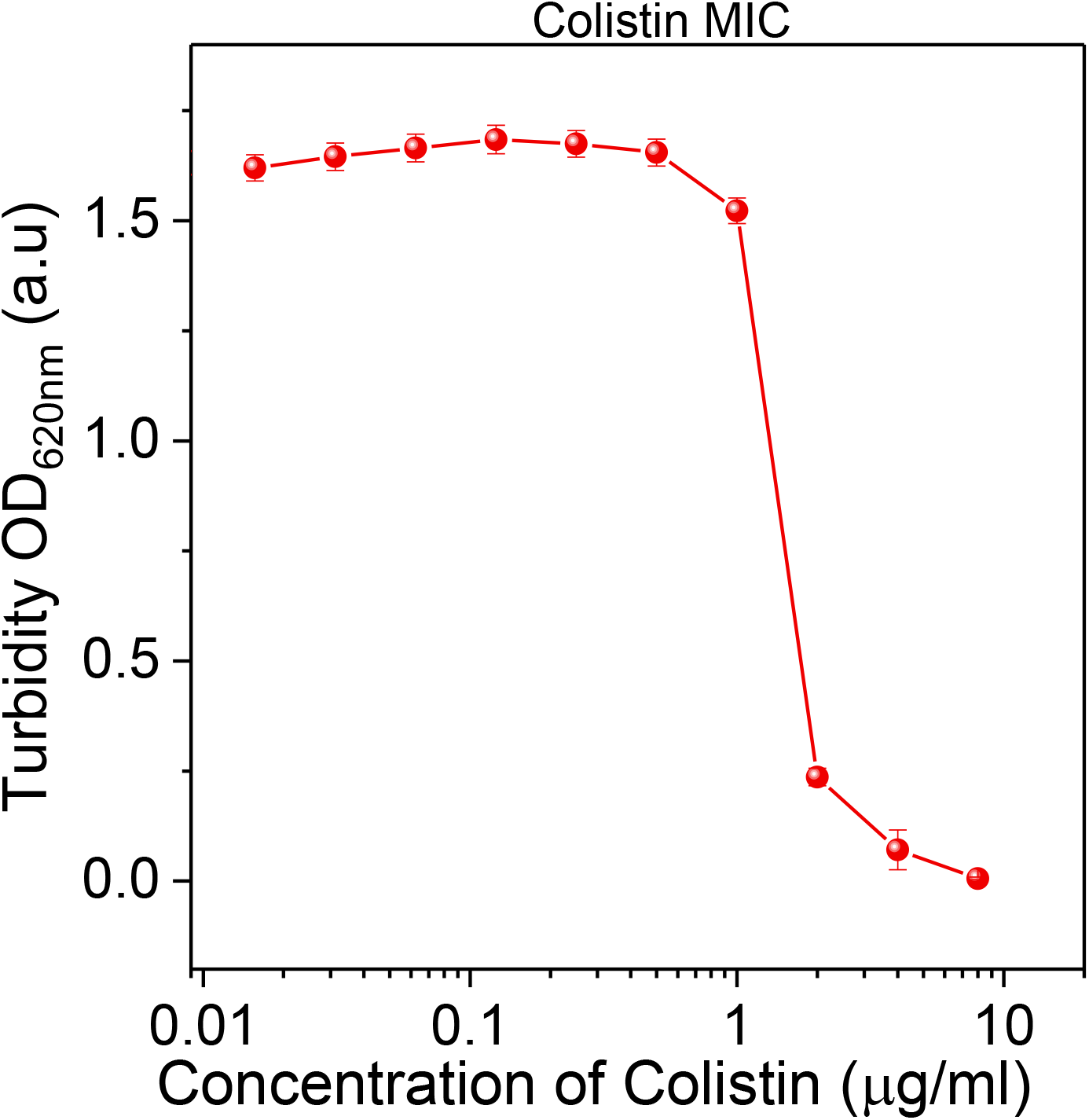
Minimum inhibitory concentration of colistin on *E. coli*: Absorbance or turbidity measurements at OD_620*nm*_ was measured as a marker for bacterial cell growth in varying concentrations of colistin to identify the minimum inhibitory concentration (MIC). For calculating the MIC of colistin, 1% primary inoculum of *E. coli* K12 was added to the growth medium with increasing concentration of colistin in a 24-well plate. It was incubated at 37 °C overnight (12 - 14 hours) with continuous shaking. The measured values indicate that the *E.coli* K12 strain has a MIC of 2 *μ*g/ml colistin. Based on this measured values, 0.25 MIC, the sub-MIC concentration used in the experiments indicate a colistin concentration of 0.5 *μ*g/ml.

**Figure S3:**
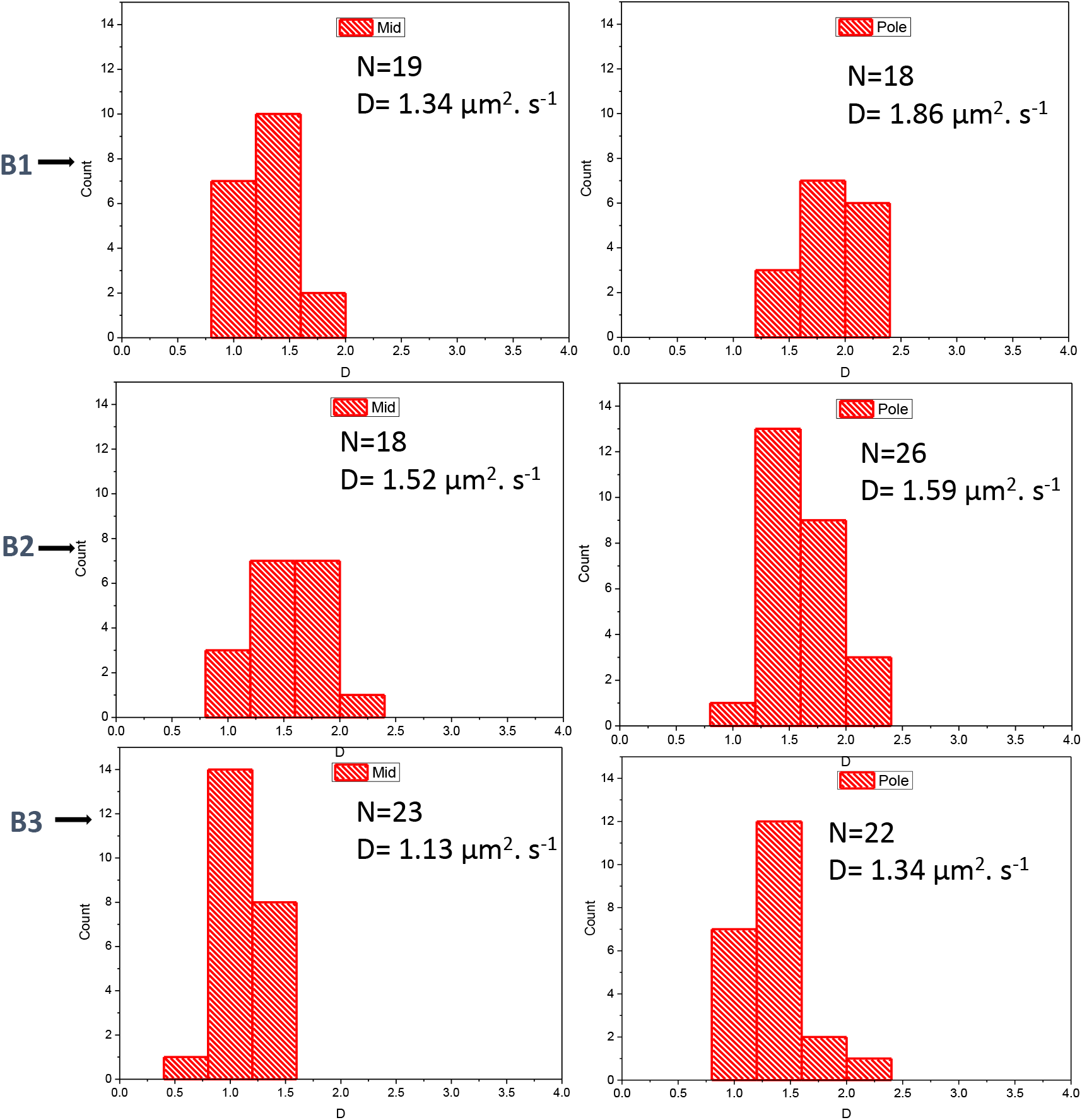
Position dependent Nile red diffusion: The fluorescence correlation spectroscopy (FCS) measurements were carried out on the polar ends and center of the membrane of untreated *E.coli* cells labelled with Nile red. The diffusion coefficients measured for three different bacteria (**B1, B2, B3**) on the center (left panels) and the polar regions (right panels) of the cell envelope are plotted as a histogram. Sample size and the mean diffusion coefficient values are provided inside the plot. Comparing the left and right panels provided we could observe that the diffusion coefficients are not significantly different from each other. Hence, lipid diffusion dynamics is not position dependent. If we compare the B1 data with B2 and B3, We can also interpret that the cell to cell variability in the lipid dynamics is also insignificant. Lipid dynamics is not dependent on individual bacterial cell and is similar in nature across the entire surface of the bacterial membrane.

**Figure S4:**
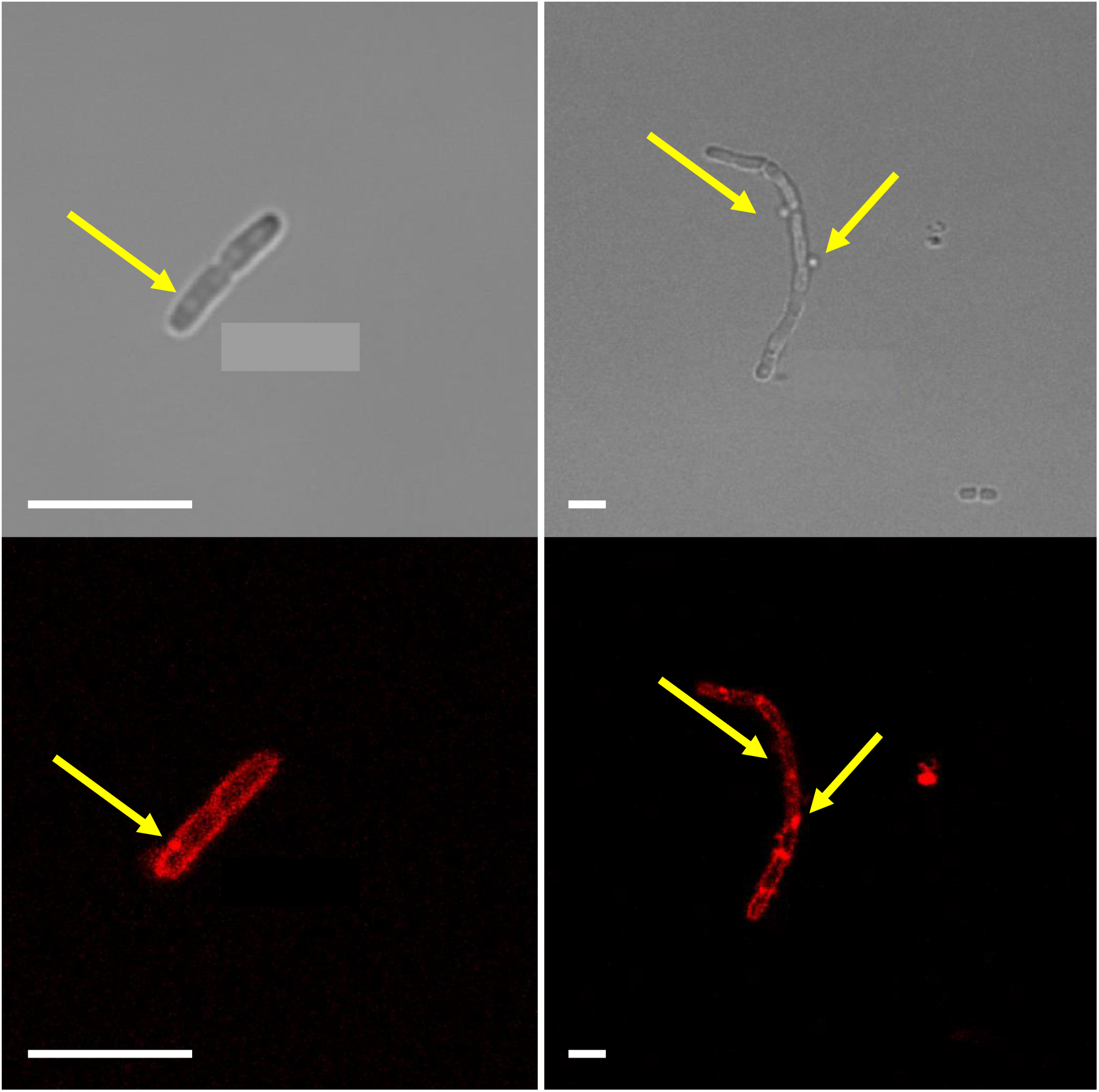
Blebs observed in colistin treated *E. coli*: Phase contrast and confocal fluorescent imaging of *E.coli* K12 cells labelled with Nile red revealed rod shaped bacterial population of uniform size and shape. When the cells were subjected to 0.25 MIC colistin treatment for 2 hours, morphological changes were observed. Cells were elongated with a significant increase in the average cell length similar to filamentation. Additionally, in few cells, small spherical bleb-like structure was observed as indicated by yellow arrows in the images. These blebs were of different sizes with enhanced Nile red fluorescent intensity. One of the hypothetical reasoning behind the formation of such blebs is that these blebs might have been formed by the bacteria as a defence mechanism by isolating the colistin throgh exocytosis pathway. However, this has to be validated in detailed specific experiments and is beyond the scope of our article. Scale bar is 4 *μ*m.

**Figure S5:**
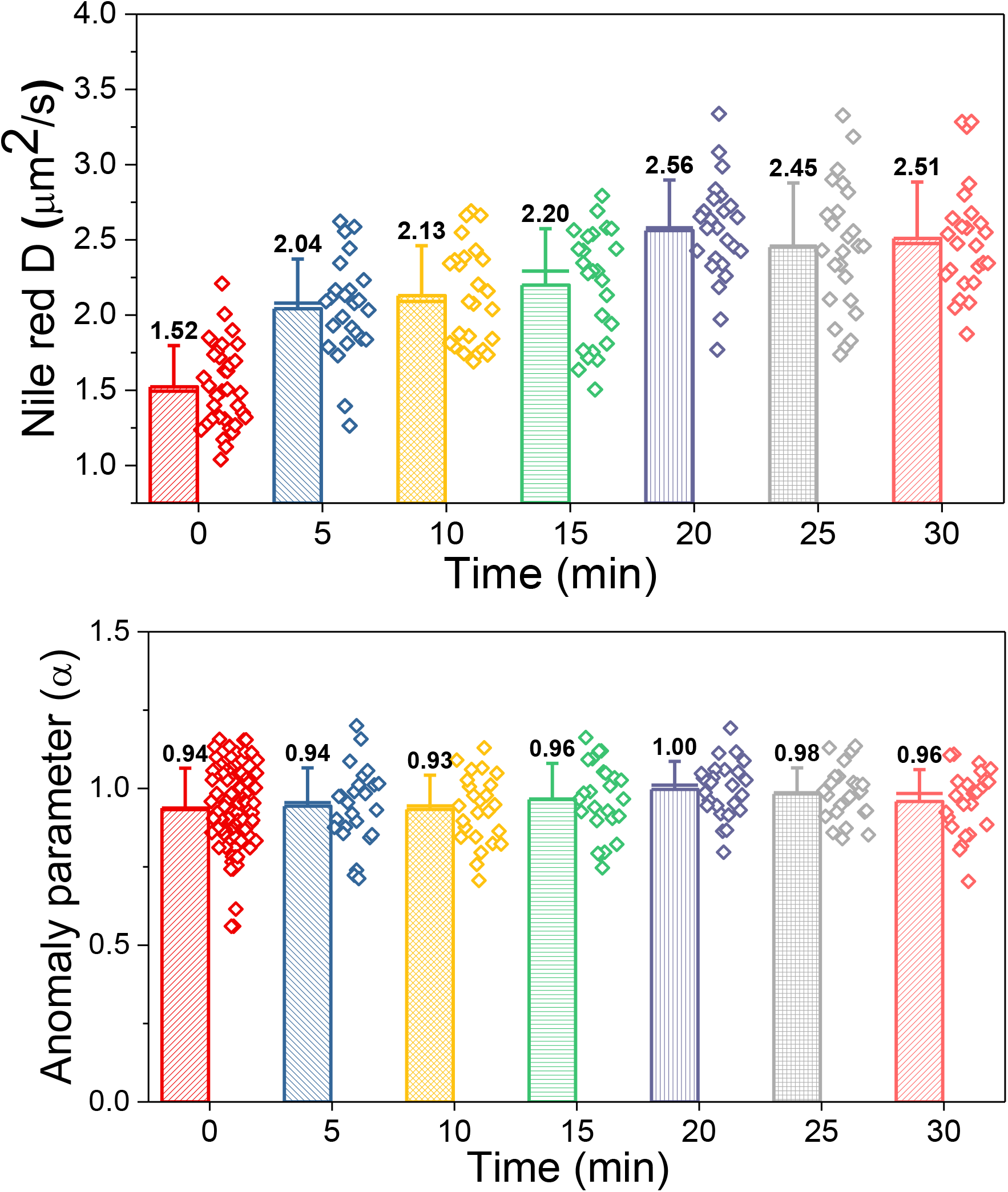
Time-dependent diffusion coefficients on colistin treatment: Inorder to identify the direct interaction of colistin on the cell envelope with respect to altered lipid dynamics, time-dependent FCS was carried out on the cells after addition of 1 MIC (1*μ*g/ml) colistin. The measured values are plotted as a function of time with the mean diffusion coefficient (top panel) and mean anomaly parameters (bottom panel) are labelled on top for each time window. The gradual increase in the diffusion coefficients observed till 20 min indicate that colistin directly interacts with the cell membrane thereby enhancing the lipid dynamics. However, the nature of the diffusion dynamics remain constant as indicated by the anomaly parameter *α*.

